# Differences in spore size and atmospheric survival shape stark contrasts in the dispersal dynamics of two closely related fungal pathogens

**DOI:** 10.1101/2023.02.08.527725

**Authors:** Jacob Golan, Daniele Lagomarsino Oneto, Shunping Ding, Reinhard Kessenich, Melvin Sandler, Tomás A. Rush, Daniel Levitis, Amanda Gevens, Agnese Seminara, Anne Pringle

## Abstract

A frequently ignored but critical aspect of microbial dispersal is survival in the atmosphere. We exposed spores of two closely related, morphologically dissimilar, and economically important fungal pathogens to typical atmospheric environments and modeled their movement in the troposphere. We first measured the mortality of *Alternaria solani* and *A. alternata* conidia exposed to ranges of solar radiation, relative humidity, and temperature. We then measured survival in an advantageous environment over 12 days. *A. solani* conidia are nearly 10 times larger than *A. alternata* conidia and most die after 24 hours. By contrast, over half of *A. alternata* conidia remained viable at 12 days. The greater viability of the smaller spores is counterintuitive as larger spores are assumed to be more durable. To elucidate the consequences of survival rates for dispersal, we deployed models of atmospheric spore movement across North American. We predict 99% of the larger *A. solani* conidia settle within 24 hours, with a maximum dispersal distance of 100 km. By contrast, most *A. alternata* conidia remain airborne for more than 12 days and long-distance dispersal is possible, e.g., from Wisconsin to the Atlantic Ocean. We observe that the larger conidia of *A. solani* survive poorly but also land sooner and move over shorter distances as compared to the smaller conidia of *A. alternata*. Our data relating larger spore size to poorer survival in the atmosphere and shorter distances travelled likely translate to other fungal species and highlight the potential for starkly different dispersal dynamics among even closely related fungi.

## INTRODUCTION

A frequently ignored but critical aspect of microbial dispersal is survival during travel. Fungal dispersal is mediated by spores, and in some species, spores are reported to cross continents or oceans in air currents (1; 2; 3). But whether spores remain viable after continental or oceanic crossings is unclear (4). As a result, an understanding of effective dispersal (defined as the fraction of spores returning to ground alive) remains elusive (4; 5). Measuring not only how far spores travel (i.e., their dispersal kernel) but also how long spores remain viable in the atmosphere (i.e., their “survival kernel”) is crucial. Tracking spores and measuring germination in nature is difficult (4; 6; 7) but measuring survival in the laboratory and connecting survival data to realistic models of movement offers one path to estimate effective dispersal.

Spore survival is often measured in terms of “germinability,” defined as the proportion of spores germinating after exposure to environmental or experimental conditions (8). Studies measuring germinability in contexts relevant to the atmosphere suggest survival is most impacted by water loss and damage from solar radiation (8; 9; 10; 11; 12). Desiccation sensitivity varies among species (13; 14) and appears to be determined by spore wall thickness, spore surface area, and relative water content within a spore (12; 15; 16; 17). High humidity is generally associated with increased germinability (8) but some species’ spores—including smut teliospores, and *Aspergillus fumigatus* and *Penicillium* spp. conidia—are released when environments are dry (10; 18; 19), perhaps to postpone germination until after deposition.

Temperature also influences germination, but temperature’s influence is not the same for every species: while colder temperatures (between 12.5 and 15.8°C) appear to maintain the germinability of *Pseudogymnoascus destructans* conidia (20), between 90-99% of *Phakopsora pachyrhizi* urediniospores fail to germinate after exposure to similarly cold temperatures (21; 22; 23). Temperature appears to be a minor influence for other species; *A. fumigatus* ascospores survive a broad range of temperatures, including heating at 70°C for 30 minutes (**24**). Some species can withstand extreme temperatures, e.g., 15% of *Cladosporium cladosporioides* conidia germinate after transient exposure to 300°C (25).

High-frequency solar radiation also influences spores’ survival (26; 27). Light in the ultraviolet (UV) spectrum (400-100nm) damages the DNA of many organisms, including fungi (9; 10; 28). Spores traveling in the troposphere are exposed exclusively to UVA (400-315nm) and UVB (315-280nm) because ozone filters shorter wavelengths (below 280nm; 29; 30). UV radiation varies significantly by latitude and altitude, and exposure changes according to cloud cover, time of day, season, and the integrity of the ozone layer at any given location (31). A spore in the atmosphere encounters variability in terms of both wavelength and dosage rate (or irradiance: W/m^2^). Some species are less resilient to UV damage (e.g., *Cladosporium herbarum*; 32) than others (e.g., *Mycosphaerella fijiensis*; 33), and other species have adapted to avoid damage, e.g., through spore melanization (*Aspergillus niger*; 34) or spore clumping (*Phakopsora pachyrhizi*; 35; 36).

A spore’s exposure to adverse humidity, temperature and solar radiation during aerial dispersal is shaped primarily by the interplay between air turbulence and gravity; these forces keep spores aloft for different times as a function of spore shape, size, or other aspects of morphology (5; 37; 38; 39; 40). Natural selection can affect potential flight times, e.g., by altering spore aerodynamics or the timing of spore release (5; 41). Fungi have also evolved traits to minimize damage from water loss or UV exposure and to navigate myriad other constraints related to movement (4; 12; 23; 38; 42; 43).

To elucidate how patterns of spore survival define the distances reached by living spores, we tested how laboratory environments relevant to atmospheric travel impact germinability. Experiments were conducted using conidia of two economically important plant pathogens: *Alternaria alternata* and *A. solani*, whose conidia and natural histories are strikingly different. While *A. alternata* is a ubiquitous, cosmopolitan species with small spores (forming chains of obovate-obtuse conidia, 10-15μm in length), *A. solani* spores are large (forming solitary, obovate-oblong conidia 75-100μm in length) and the species is primarily associated with solanaceous (especially potato and tomato) crops (44; 45; 46; 47). Both species pose serious threats to solanaceous crops and conidia often co-infect the same plant (47; 48).

In a first experiment (Experiment 1), we exposed conidia of *A. alternata* and *A. solani* to a range of relative humidities (RH), temperatures (T), and UV wavelengths and intensities (UV) for 96 hours. Data were used to identify combinations of RH, T and UV favorable to the retention of germinability. In a second experiment (Experiment 2), we exposed approximately 10.0^6^ and 10.5^5^ spores of *A. alternata* and *A. solani* to a favorable environment for over 12 days (288 hours), a timescale relevant to continental or oceanic dispersal (1; 2; 49; 50). We next used simulations of particle transport in atmospheres to model the dispersal of spores (51; 52). Ultimately, patterns of effective dispersal emerge as strikingly different between these two closely related species.

## MATERIALS & METHODS

### Overview

In Experiment 1 we exposed conidia of *A. alternata* and *A. solani* to open air with different combinations of ultraviolet wavelengths and irradiance (UV), relative humidities (RH), and temperatures (T). We chose RH and T ranges relevant to spores dispersing in the troposphere and tested ten combinations (1-10, Table S1) typical of central Wisconsin in summer (53; 54). We conducted experiments in a single controlled environmental chamber at the University of Wisconsin Biotron (Madison, WI, USA) and the ten combinations of RH-T were tested sequentially in this single chamber. For each RH-T combination, we tested 21 UV strengths, including both realistic and unrealistic irradiances (56; 57; 58; Table S2) for a total of 10 RH-T conditions x 21 UV strengths = 210 treatments per species. We ran each iteration of Experiment 1 for 96 hours. We measured germinability at 24, 48, 72, and 96 hours. Next, we sought to understand how long conidia could live in a nearly ideal environment, an experiment designed to test the maximum potential reach of each species. In Experiment 2 we used a combination of UV-RH-T favorable to the retention of germinability as a single environment in two experimental runs, one for *A. alternata* (2A) and a second for *A. solani* (2S). We conducted Experiment 2 for 288 hours (12 days) and measured germinability at 0, 24, 48, 72, 144, 216, and 288 hours (or days 0, 1, 3, 6, 9, and 12). Methods used to collect and generate *A. alternata* and *A. solani* conidia are found in Supporting Information 1.

### Exposing spores to different combinations of UV-RH-T (Experiments 1 and 2)

*Physical setup:* For Experiment 1, a series of plexiglass platforms were cut and fit as steps into a frame made of PVC pipes (Figure 1). Because irradiance is inversely proportional to the squared distance between a light source and a surface, each graduated step was exposed to a different intensity of UV (compare Figure 1 and Table S2). Each plexiglass step measured 20.32 cm wide by 66.04 cm long; six steps were placed under 40 W_UVA_, six under 40 W_UVB_, four under 15 W_UVA_, four under 15 W_UVB_, and one in 0 W (i.e., complete darkness); a total of 21 steps or surfaces.

**Figure 1:**
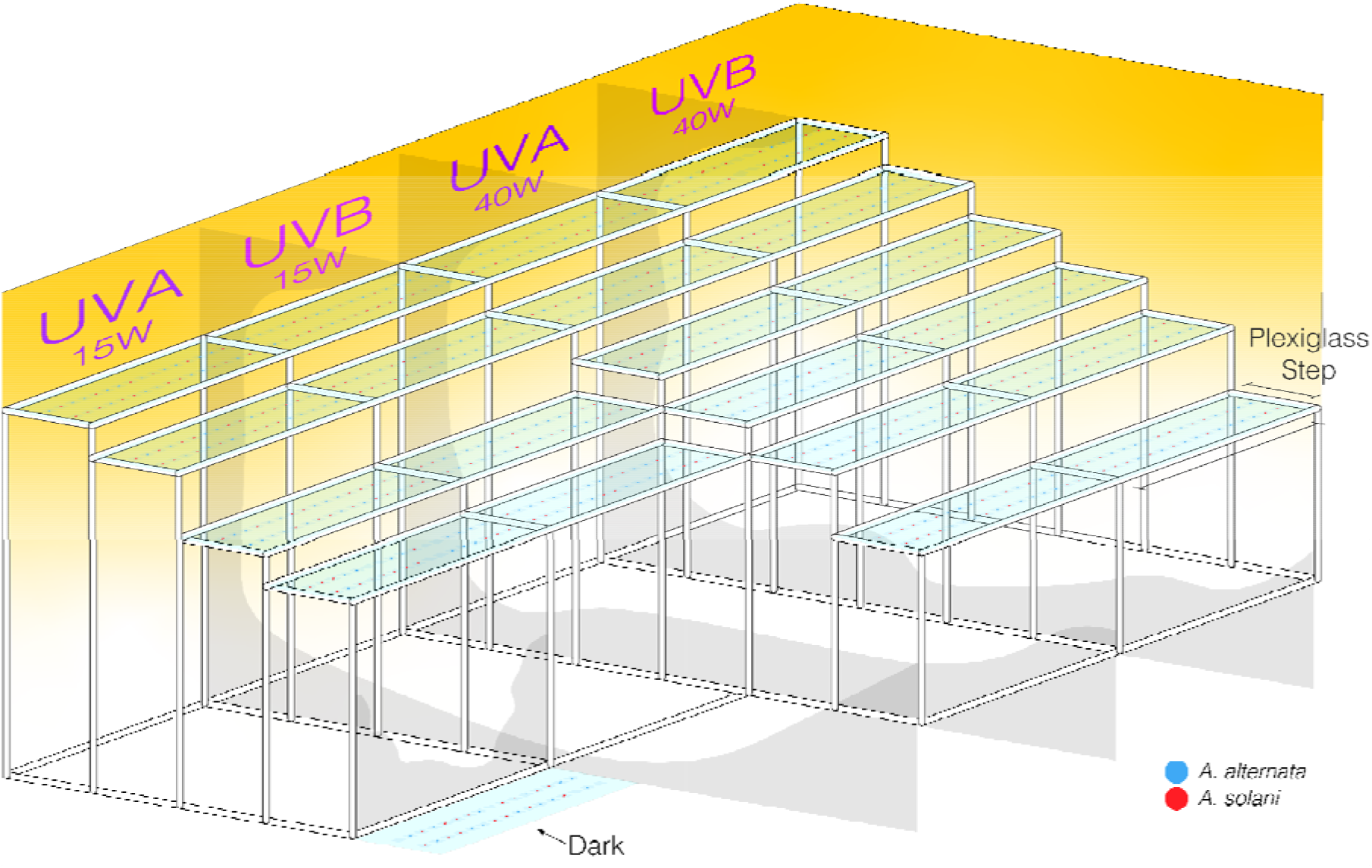
Experiment 1 experimental setup. Clear plexiglass surfaces (or “steps”) were arranged at different heights beneath a UVA or UVB light source. Quartz cover slips coated in spore suspensions of either *A. alternata* or *A. solani* were placed on each plexiglass step using a randomized block design. UV bulbs were suspended above the 20 steps (an additional step was kept in complete darkness, labelled “Dark”). A module encompasses all surfaces underneath one of the four UV bulbs of a given wavelength (UVA or UVB) and power (Watts). There are four modules. The experimental apparatus was placed in an environmental chamber and a single relative humidity (RH) and temperature (T) specific to one of ten environments (conditions 1-10) was maintained and monitored every five minutes by an automated system during each of the experimental runs (1-10). Experimental runs took place in series, one after the other, using the same chamber. Lights cycled through a 12-hour on, 12-hour off schedule for the 96 hours of each experimental run and four cover slips were sampled from each step at 24-hour intervals. Black plastic tarp was placed between each module to prevent leakage of UV light between modules. Conidia did not germinate on cover slips.

For Experiment 2, a single 121.92 cm (48 in) long by 66.04 cm (26 in) wide plexiglass step or platform was placed under a light source at a strength consistent with the single treatment chosen from Experiment 1.

#### Conidial manipulation

Experiment 1 conidia were first placed on microscope coverslips. Coverslips were prepared by spreading 50 μL of a gently mixed, concentrated conidial suspension onto the upper surface of a sterile 19×19 mm ultra-thin (0.25mm) quartz cover slip (Chemglass Life Sciences, Vineland, New Jersey, USA). Coverslips were left to dry in darkness for a few minutes before being placed in the environmental chamber. For each of Experiment 1’s 10 conditions, coverslips were placed as two rows of 16 on each step (32 coverslips per UV-RH-T treatment; 16 for each species, Figure 1); coverslips were arranged according to a randomized block design. As each of the 10 conditions included a total of 32 coverslips for each of 21 treatments the total number of coverslips for each experimental run was 672 (336 coverslips per species). In total, the 10 conditions involved 6,720 coverslips.

Experiment 2 conidia were spread onto glass slides instead of coverslips. A total of 238 25×75 mm glass microscope slides (Globe Scientific, Mahwah, New Jersey, USA) per species were coated in 200 μL of conidia suspensions and left to dry in darkness for a few minutes before being placed in the environmental chamber. For each of the two runs (2A and 2S), a total of 217 slides were randomly placed as a grid across the single plexiglass platform. The remaining 21 slides were kept in complete darkness.

#### Light treatments

In Experiment 1, UVP XX-Series UV Bench Lamps (Analytikjena, Jena, Germany) were suspended above the plexiglass steps (Figure 1) to generate different intensities of UV (Table S2). Irradiances were measured for each step with a UV Light Meter (Sper Scientific Direct, Scottsdale, Arizona, USA) at the start of each experimental run (Table S2). To prevent leakage of UV light from one module to another, black plastic fabric was placed between modules, and the UV Light Meter was used to confirm both that no light was leaking between modules and that the step kept in darkness was dark. In Experiment 2, fixtures emitting only UVA (6.29±0.17 W/m^2^ for both species) were placed above the single treatment surface. In both experiments, day-night cycles were approximated by alternating 12 hours of continuous UV irradiation with 12 hours of darkness.

#### Relative humidity and temperature

In Experiment 1, the environmental chamber was calibrated to one of the 10 RH-T conditions (Table S1). These RH and T values are typical of central Wisconsin during the peak seasonal concentrations of airborne conidia of A. alternata and A. solani (54; 55). In Experiment 2, a single RH and T found to favor the retention of germinability for *A. alternata* (RH=90%, T= 15°C) and *A. solani* (RH=90%, T= 20°C) was held for 288 hours. In both Experiments 1 and 2, RH and T were monitored every five minutes to ensure conidia were consistently exposed to a given treatment.

### Measuring germinability

#### Imaging

Conidia were germinated according to methods provided in Supporting Information. After 24 hours conidia were counted (*N_total_*). The slide holder on an Olympus CX31 compound microscope (Olympus, Tokyo, Japan) was removed so that conidia could be observed directly from agar plates. All conidia were visualized using an Olympus PlanApo N 2x objective lens (Olympus, Tokyo, Japan). To increase light penetration through agar, the microscope light condenser was removed. Digital images were captured using a Canon EOS Rebel II (Canon, Tokyo, Japan) with a Martin Widefield 1.38x DSLR adapter for Olympus BX and SZX with 51 mm dovetail photoport (Easley, South Carolina, USA), resulting in a total magnification of 2.76x. In Experiment 1, ten non-overlapping images of conidia were randomly captured from each plate at each condition and time, and the number of germinated spores was counted (*N_germinate_*).

In Experiment 2 the same protocols were followed but five images were captured per plate for *A. alternata* and 20 images were captured per plate for *A solani.* Image numbers differ to account for differences in the density of conidia observed between species.

#### Image processing

Custom algorithms developed by MIPAR v3.2 (Worthington, Ohio, USA) were used to count germinated and ungerminated conidia. Conidia size and germ tube development are different for the two *Alternaria* species, and as a result, species-tailored counting algorithms were used. A full description of image processing protocols is found in Supporting Information 2. In brief: out-of-focus features of each image were removed, as were features outside of the size range of conidia. Thresholding substantially reduced noise caused by debris and uncountable clusters of conidia (Figure 2). Remaining features were then classified as either germinated or ungerminated conidia. To ground truth the counting algorithms, 50 images of *A. alternata* and *A. solani* were randomly selected and germinated and ungerminated conidia counted by eye. Manual counts of live and dead conidia were compared to results generated from our custom software (Supporting Information 1; Figure S1).

**Figure 2:**
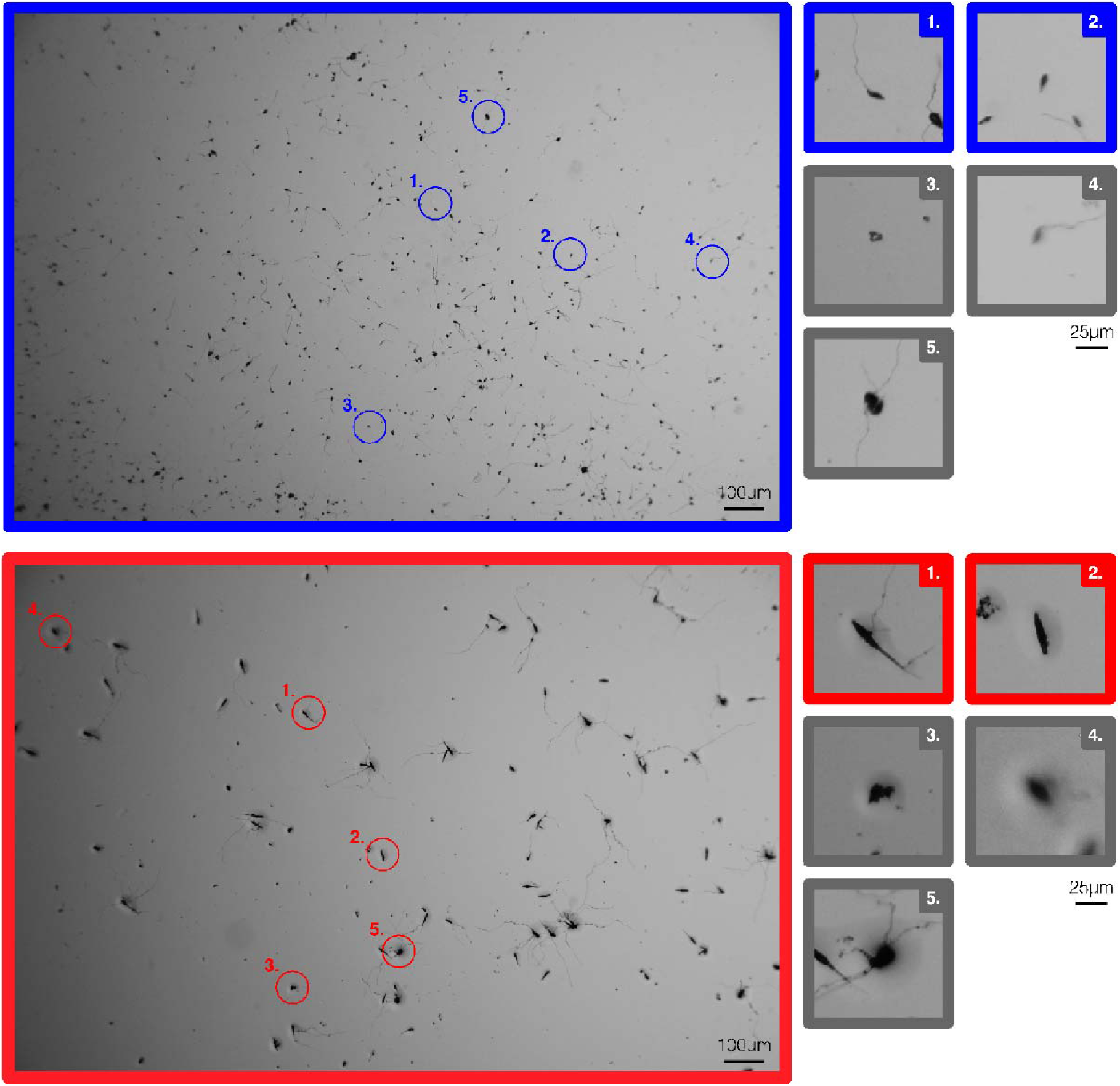
Example images of germinating conidia of *A. alternata* (blue) and *A. solani* (red) on water agar plates. Boxes 1 and 2 show germinated and ungerminated conidia, respectively. Boxes 3-5 illustrate debris, out-of-focus conidia, and uncountable clusters of conidia, respectively.

### Statistical analyses

#### Mixed effect models

We used the R package *glmmTMB* (59; 60) to test for significant differences among the numbers of germinated conidia across treatments in Experiments 1 and 2. The number of germinated and ungerminated conidia was calculated per coverslip and modeled using a log link function and log-transformed mean total number of conidia as an offset (59).

Experiment 1 data: variables included days of exposure, UV wavelength (including darkness), distance from a UV light source, RH and T; each species was analyzed separately. Random effects were included to account for any deviations in environmental chamber performance or for fluctuations in UV intensity across a step (Figure 1). We computed full models and then simplified models by removing uninformative variables using the corrected Akaike Information Criterion (AICc) (Table 1). In addition, we performed Tukey’s *post-hoc* tests to correct for multiple comparisons of means (Table S3(B); 60).

**Table 1.**
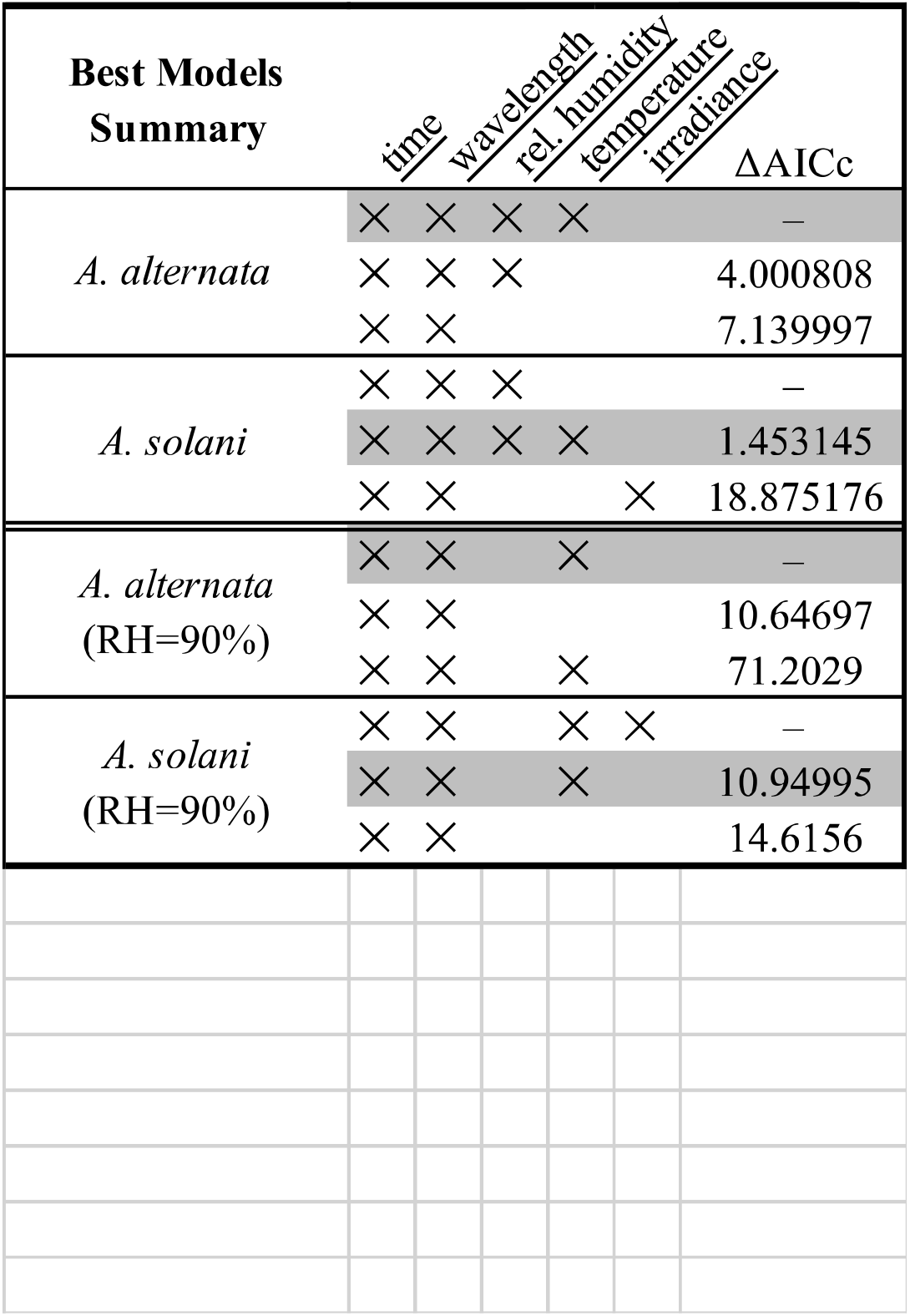
A. Best model summaries for Experiment I conditions 1-10. B. Summary of multiple comparisons of means using Tukey contrasts for best fit mixed models. Contrasts are shown for A. alternata (blue), and A. solani (red) for relative humidity (lower triangle) and temperature (upper triangle). Main values are pairwise estimates and values in parenthesis are p-values.

#### An additional analysis

In a separate analysis of Experiment 1 data, we tested for significant effects of UV, RH and T on conidia germination using Kruskal–Wallis tests followed by *post-hoc* assessments of significance using Dunn’s multiple comparisons with a Benjamini-Hochberg adjustment (Table S4; R Core Team 2020). Germination at hour 96 only was compared across (a) UV wavelengths (UVA, UVB, and darkness), (b) RHs per UV wavelength (e.g., UVA-50% RH vs. UVA-90% RH), (c) Ts per UV wavelength (e.g., UVB-20°C vs. UVB-15°C), and (d) conditions (e.g., Experiment 1 condition 1 vs. Experiment 1 condition 2, etc.).

### Using models of atmospheric transport to simulate dispersal across space over time

To understand how patterns of germinability affect the movement of both *Alternaria* species across North America, we modelled the transport of *A. alternata* and *A. solani* conidia in the atmosphere. A full description of model parameters and methods is found in (5). Briefly: numerical simulations tracked many representative trajectories of spores in the atmosphere using meteorological data available from the National Oceanic and Atmospheric Administration (NOAA) and the Hybrid Single-Particle Lagrangian Integrated Trajectory (HYSPLIT) model (61). Specifically, we used the North American Regional Reanalysis described in (62), as it combines numerical simulations with observational data.

The movement of conidia through the atmosphere was modeled vertically and horizontally, with gravitational settling velocities proportionate to conidial dimensions: 20×7.5μm for *A. alternata* and 100×10μm for *A. solani* (spore density approximated as 1 g/cm^3^; 63). Models simulate dry deposition by randomly removing spores that travel close to the ground using a constant rate proportional to the deposition velocity. Turbulent eddy diffusivity was estimated following Beljaars & Holtslag (“BH”; 64).

In each simulation, a total of 500 000 conidia of each species were released from central Wisconsin (44.119N, −89.536W) at an altitude of 10 m. Simulations were run per species with the following initial conditions: July 15, August 1, August 15, and September 1 at 0:00, 10:00, and 14:00 hours for the years 2009-2018 (a total of 240 combinations of parameters). Dates and times were chosen based on historical data of peak conidial concentrations (54). Simulations lasted 288 hours, at which point the latitude, longitude, maximum height, and time of deposition were recorded for each of the 500 000 conidia released at time zero per each of the 240 parameter sets.

The output of each simulation was imported into R v3.6.2 (R Core Team). The distance travelled by each spore from take-off to deposition was calculated using the WGS84 terrestrial reference system with *geosphere* v. 1.5-10 (65). To visualize the geographic spread of conidia, data were aggregated by date of release and year. Landing times were grouped into six-hour intervals from zero to 266 hours. The centroid of the spatial range traveled by all conidia within each six-hour interval was calculated and an ellipse was drawn around each centroid, with the major axis oriented in the direction of maximum spread from the centroid. The major axis radius is equal to the standard deviation of the distance travelled along the direction of the major axis by spores that sediment within the six-hour interval. Similarly, the minor axis represents one standard deviation of the distance travelled in the direction perpendicular to the major axis by all spores that sediment within the 6-hour interval. Ellipses were calculated using *aspace* v3.2 and custom in-house scripts (https://github.com/jacobgolan/Alternaria_Dispersal.git; 66). To minimize two dimensional distortions of spore trajectories across Earth’s curved surface, the R package *sp* v.1.4-0 was used to correct the latitude and longitude of each spore from an EPSG:2288 coordinate system to EPGS:4326 (67).

## RESULTS

### Counting germinated spores

Germinability was successfully quantified for *A. alternata* and *A. solani* conidia using automated counting algorithms: automated and manual counts are strongly correlated (Figure S2).

### Identifying parameters most likely to maximize spore germination (Experiment 1)

#### Fitting models

Experiment 1 data enabled identification of the combinations of UV, RH and T resulting in greatest numbers of germinated conidia (Table 1). Full models were computed using time, wavelength (UVA, UVB or darkness), RH, T, and irradiance (W/m^2^; Figure 1) and simplified final models were chosen by comparing models’ Akaike Information Criterion (AIC).

The number of germinated conidia on each coverslip was modeled as a random variable distributed according to a negative binomial distribution. The expected value of the distribution, conditioned on each treatment, took the form:

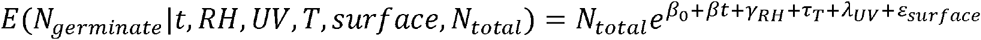

where *N_total_* is the total number of conidia on a coverslip (alive and dead); *β* is a parameter quantifying how quickly germination decreases and *t* is time of exposure to a specific condition (in days); *γ_RH_*, *τ_T_* and *λ_UV_* are parameters quantifying the effects of RH, T and exposure to UV light. *ε_surface_* represents the random effects on each surface (a random variable distributed according to a Gaussian centered at zero and with a standard deviation *σ*). The fit produces estimates for our nine coefficients of interest *β*, *γ*_60%_, *γ*_75%_, *γ*_90%_, *τ_15°C_*, *τ_20°C_*, *τ_25°C_*, *λ_UVA_* and *λ_UVB_*) and we choose RH = 50%, T = 10°C and no UV exposure as a reference condition, hence *γ_50%_ = τ_10°C_ = λ_dark_ = 0*. Exponentiated coefficients greater than 1 translate to an increase in germinability with respect to the reference condition, and exponentiated coefficients less than 1 translate to a decrease in germinability with respect to the reference condition (Figure 4). *β*_0_ is the intercept accounting for dead spores in the reference condition at *t* = 0.

**Figure 3:**
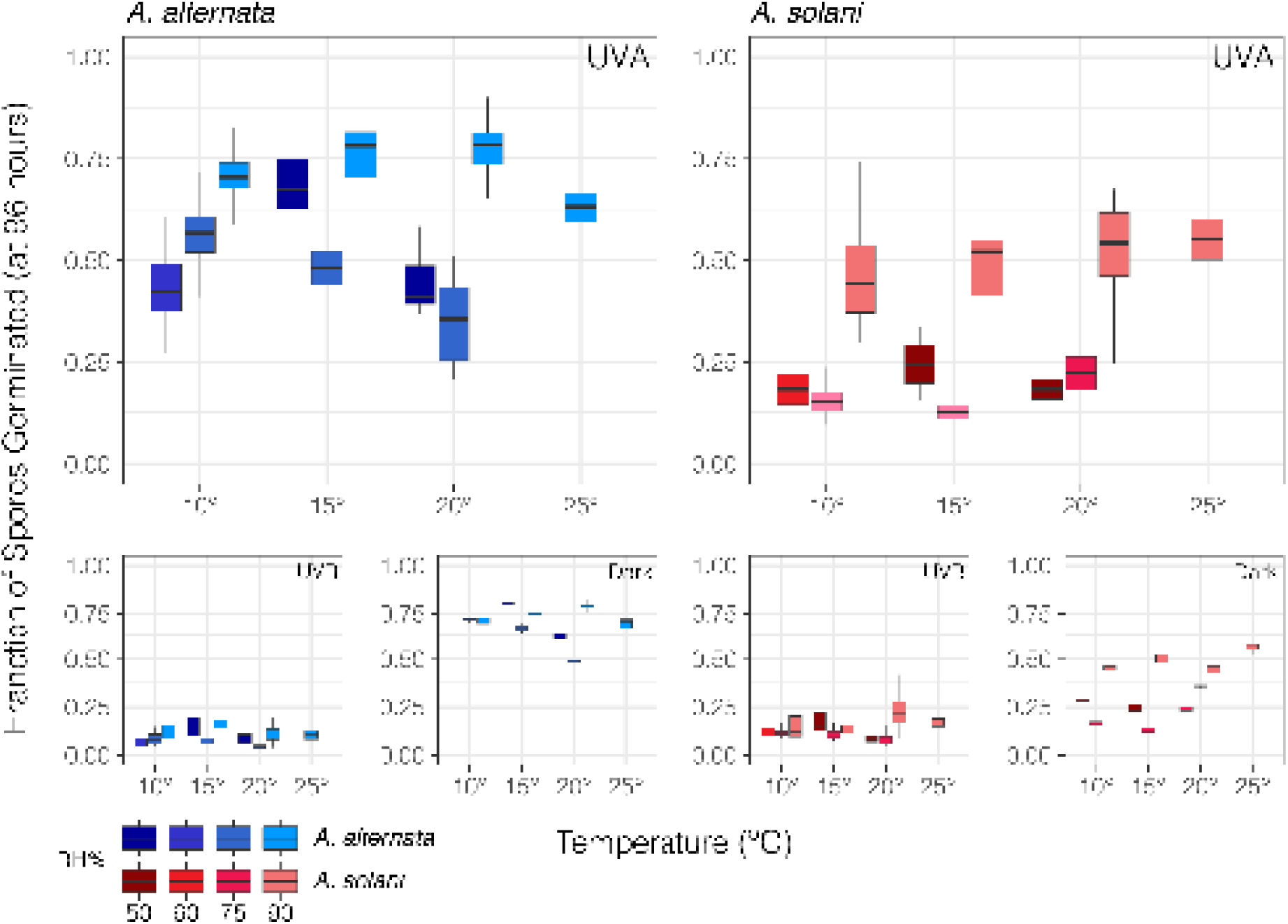
Experiment 1: Proportion of conidia relative to initial number of conidia germinating after 96 hours for all tested relative humidities (RH %), temperatures (T °C), and UV dosages. Data of *A. alternata* are blue and data of *A. solani* red. Tones of blue and red mark different RH environments.

**Figure 4:**
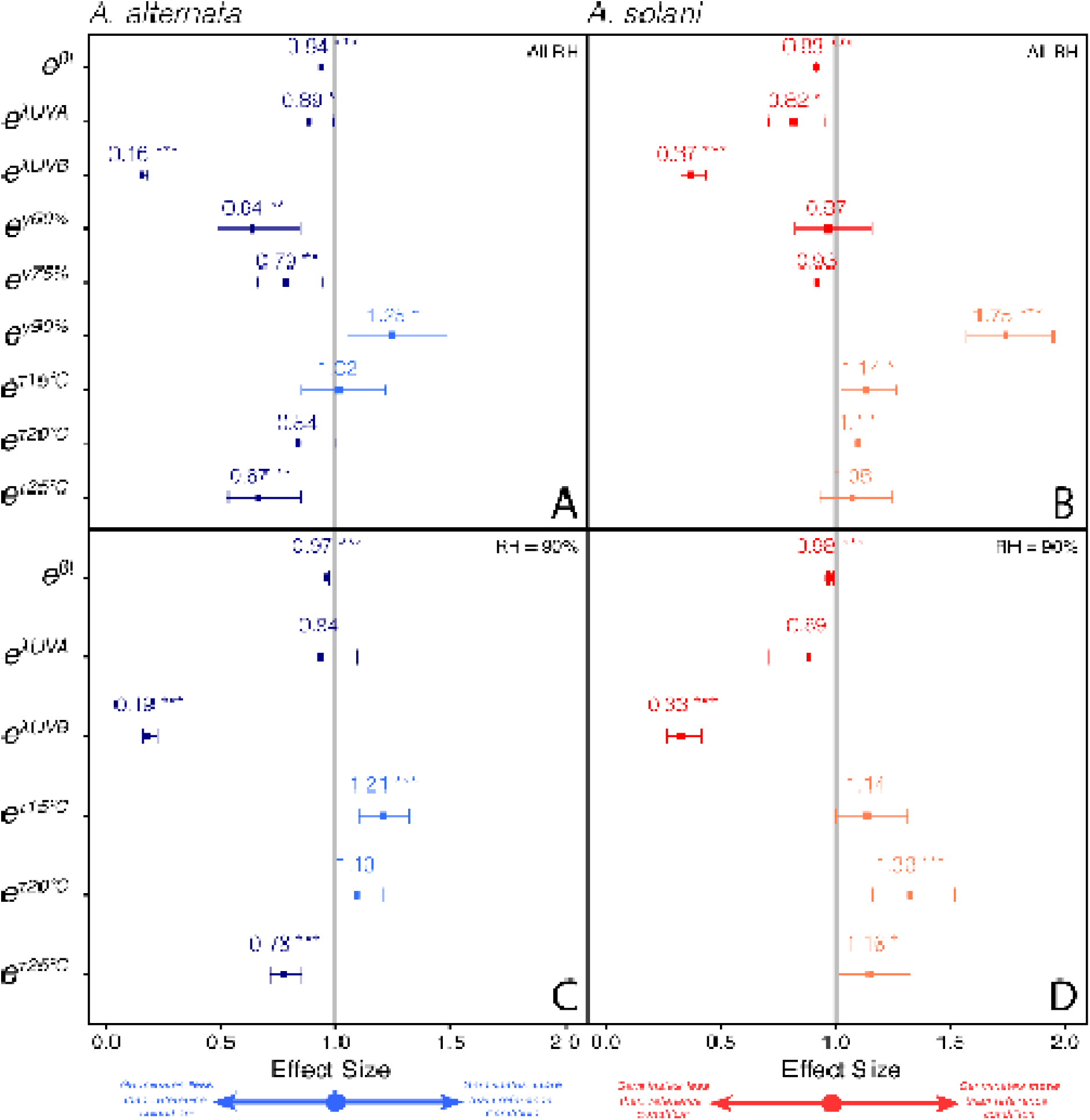
Summary of effect sizes for parameters included in best-fit generalized linear mixed models. Effect sizes shown in exponentiated form. Panels A) and B) show estimates for models including all ten Experiment 1 conditions, and (C) and (D) show effect sizes from models fitting only conditions 7-10, or the conditions for which RH was held at 90%. Values for *A. alternata* and *A. solani* shown in blue and red, respectively.

We next compared models’ AIC to identify the minimum number of parameters needed to explain experimental data without overfitting. The best-fitting model of *A. solani* conidia germination did not include T, but to enable comparisons between *A. solani* and *A. alternata,* we selected the second-best *A. solani* model, which included T and was identical to the best fit *A. alternata* model (Table 1).

Models identify both UV wavelengths as detrimental to germination (*e^λUVA^* = 0.89 and 0.82, and *e^λUVB^* = 0.16 and 0.37, for *A. alternata* and *A. solani* respectively, Figure 4). Conidia kept in darkness germinated most readily and UVB exposure resulted in the smallest numbers of germinated conidia (Figure 3). While we observed differences in conidial germinability among different wavelengths (Figure 3), selected models did not include irradiance (W/m^2^) as a parameter (Table 1). Kruskal-Wallis followed by *post-hoc* Dunn tests confirm this result (Table 4).

Relative humidities of 90% maximized germination at all temperatures and UV wavelengths (*e^γ90%^* = 1.25 and 1.75 for *A. alternata* and *A. solani* respectively, Figure 4). Kruskal-Wallis followed by *post-hoc* Dunn tests confirm this result (Kruskal-Wallis *χ*^2^ = 225.05, 77.664, 28.624, respectively, all with df = 3, p-value < 0.0001 for each; Table S4).

Results for T were less consistent than results for RH or UV. Models suggest 15°C maximized germination for both species (Figure 4), but *A. alternata* conidia kept at 90% RH appear to germinate equally well at both 15°C and 20°C (p-value < 0.05; Table S3, Figure 3). Kruskal-Wallis followed by *post-hoc* Dunn tests were also inconclusive (*χ*^2^ = 11.28-55.14, df = 3, 0.01 < p-value < 6.40×10^-12^; Table S4). Because 90% RH clearly maximized the germination of both species’ conidia, temperature was reinvestigated using only the four conditions (7-10) involving 90% RH (Table 1, Figure 4):

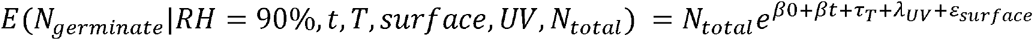

*E*(*N_germinate_*|*RH* = 90%, *t, T, Surface, UV, N_total_*) = *N_total_e*^*β*_0_+*β_t_*+*τ_T_*+*λ_UV_*+*ε_surface_*^ Results were more consistent; according to both model effect sizes (Figure 4, Table 3, Table S3), and Kruskal-Wallis and *post-hoc* Dunn tests (Table S4), 15°C is the most favorable temperature for *A. alternata* germinability, and 20°C is the most favorable for *A. solani* germinability.

Based on these results, parameters chosen for Experiment 2 included an RH of 90% and T of 15°C for *A. alternata* (2A), and 90% RH and 20°C for *A. solani* (2S). We exposed conidia to alternating periods of 12 hours UVA light and 12 hours darkness at an irradiance of 6.29±0.17 W/m^2^, equivalent to the lowest UVA-40W dosage administered in Experiment 1 and a UV environment typical of the troposphere (Table S2; **29**).

### Measuring spore germination over timescales consistent with long distance dispersal (Experiment 2)

The two *Alternaria* species demonstrated markedly different germination patterns over 288 hours. A greater total number of conidia and proportion (i.e., fraction of total conidia) of *A. alternata* conidia germinated at all sampling points (hours 0, 24, 72, 144, 214 and 288), compared to *A. solani* conidia (Figure 5). Germinability of *A. alternata* conidia decreased linearly over time, but germinability of *A. solani* conidia fell sharply within 24 hours and subsequently plateaued. Germinability remained at approximately 12-20% after 24 hours and a visual inspection of *A. solani* conidia suggests most conidia germinating after 24 hours develop atypical germ tubes, compared to conidia germinating at 0 hours (Figure S5). These abnormally growing conidia could not be measured by custom MIPAR algorithms because they were designed to provide a binary classification (germinated/ungerminated). Atypical conidia grew germ tubes reaching a length of approximately 100-150μm (compared to ~200μm or more at 0 hours) and germ tube growth was delayed (Golan pers. obs.). Differences between *A. alternata* and *A. solani* germination are corroborated by Experiment 1 data: the germinability of *A. alternata* conidia decreases linearly over time, but germinability of *A. solani* conidia falls sharply within 24 hours of the start of the experiment (Figure S6). In Experiment 2, the half-life of germinability for *A. alternata* is approximately 35 hours (i.e., ~2% loss in germinability per hour under UVA). In stark contrast, the half-life of germinability for *A. solani* is approximately 1.5 hours (i.e., ~47% loss in germinability within the first 24 hours).

**Figure 5:**
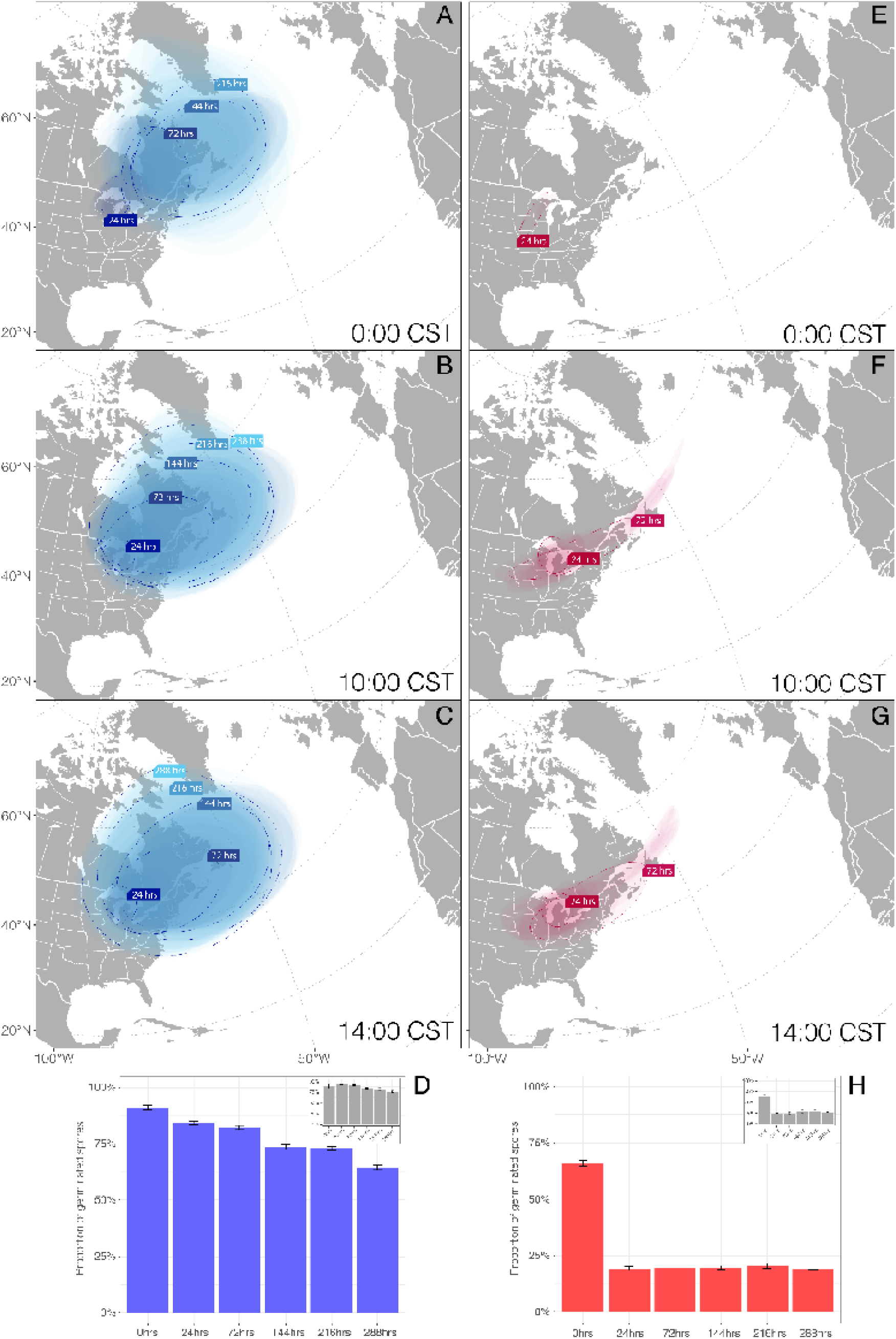
(A-C, E-G) Spatial visualization of HYSPLIT models. Maps show release times 0:00, 10:00 and 14:00 CST averaged over ten years. Simulations for *A. alternata* and *A. solani* are shown in blue and red, respectively, with colors becoming less intense as time increases. Data for each six-hour interval are summarized as ellipses with major axes is in the direction of maximum spread, and lengths as one standard deviation of the mean distance travelled in each 24-hour period (ranges marked with dotted outlines). (D & H) Proportion of germinated spores averaged per block (slide), Experiment 2. Data in blue and red for *A. alternata* and *A. solani*, respectively. Data for spores kept in darkness shown in grey inset.

#### Effective Dispersal

The HYSPLIT simulations of conidia dispersing from central Wisconsin show the smaller conidia of *A. alternata* as travelling over greater ranges than the larger conidia of *A. solani* (Figure 5; Figure 6). But spore size does not affect the speed of movement, instead, it affects the altitude of spores and controls how long a spore will remain aloft. The number of *A. solani* conidia in the air decreases two to three times faster than the number of *A. alternata* conidia in the air (Figure 6). By 144 hours (day 6) no *A. solani* conidia remain aloft (in any simulation). By contrast, at 288 hours (day 12) significant numbers of *A. alternata* conidia are still found in the atmosphere (in all simulations). Before all *A. solani* conidia settle, they can travel as far as *A. alternata* (more than ~3,000 km, Figure 6), but the number of conidia reaching these long distances is less than 1% of the total released, as compared to *A. alternata.*

**Figure 6:**
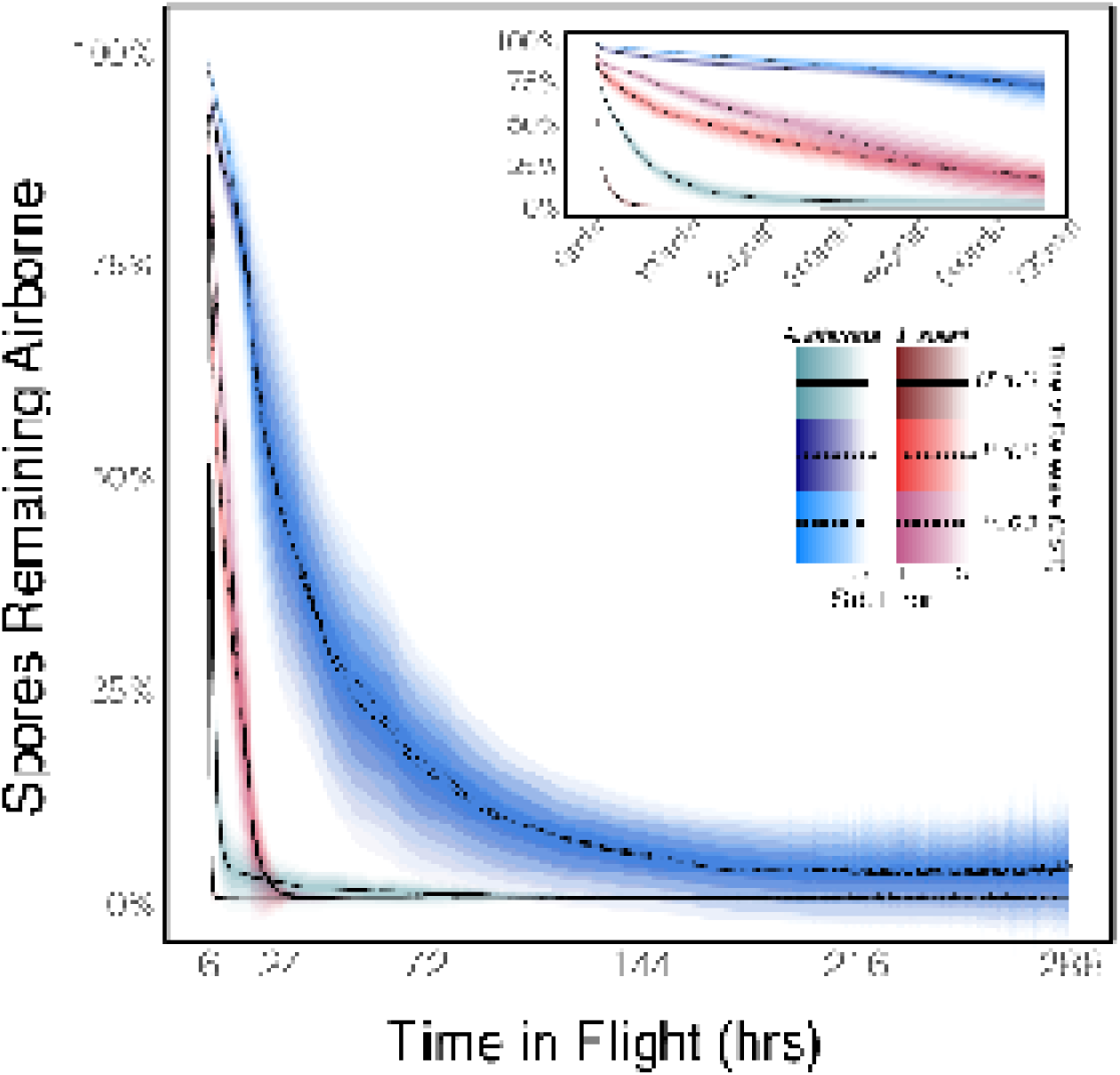
Results of HYSPLIT models of spore dispersal showing spores remaining airborne as a function of time after take-off. Small insets show the same data for the first 12 hours. Spores remaining aloft over time shown in blue and red for *A. alternata* and *A. solani*, respectively. Trajectories were simulated for three release times: 0:00 (solid line), 10:00 (dotted line), and 14:00 (dashed line). All conidia of *A. solani* settle before the end of the 288-hour simulation. Shades correspond to the number of standard errors from the mean, means represented by black trend lines.

Similarities are greatest for the two species at release times 10:00 CST and 14:00 CST, during midday when wind turbulence is greatest. At a release time of 0:00 CST, *A. solani* conidia settle to the ground before day three, while *A. alternaria* conidia are reaching Greenland on day nine. But even at release times 10:00 CST and 14:00 CST, dispersal dynamics are very different for the two species (Figure 5). Ranges for *A. solani* are elongate and rather narrow, compared to the more circular, broader ranges of *A. alternata,* and the edges of *A. solani*’s ranges will involve many fewer conidia, as compared to the edges of *A. alternata*’s ranges (Figure 6). Because the germinability of *A. solani* conidia declines rapidly most or all spores at its ranges’ edges will be inviable.

## DISCUSSION

We tested how temperature, relative humidity, UV light exposure, and their combinations affect the germinability of *A. alternata* and *A. solani* (8; 12). Next, we measured survival in a favorable environment over a timescale consistent with continental dispersal. We combined the survival data with models of spore movement to offer a realistic bound on the effective (as opposed to potential) dispersal of spores in the atmosphere (4; 5). We specifically chose to measure longer-timescale survival in a favorable and realistic, but unnaturally static, tropospheric environment to probe the edges of the potential reach of spores, asking, “how far would the ‘luckiest’ spores of either species travel”?

The effective dispersal of the two species is very different. As an illustration, consider the ability of *A. solani* and *A. alternata* to reach Maine (a potato growing state) when both are released at 10:00 CST: less than 1% of *A. solani* reach Maine and most are inviable; by contrast, upwards of 25% of *A. alternata* reach Maine and 75% of them are still viable (Figure 5; Figure 6). The combination of more time aloft and greater longevity results in a larger number of *A. alternata* conidia travelling hundreds to thousands of kilometers and landing still able to cause infection (Figure 5). Less than 1% of *A. solani* conidia are still in the atmosphere after 24 hours and because these spores either cannot germinate or germinate abnormally, they are unlikely to cause disease.

The conidia of *A. alternata* are both small enough to travel over 1 500 kilometers and physiologically equipped to survive the journey (3; 68; 69; 70). We can find no data addressing the global population biology of *A. alternata,* but based on our experiments we hypothesize it may function as a single, global population, similar to *Aspergillus fumigatus* (71).

The larger conidia of *A. solani* are more vulnerable to atmospheric hazards than the smaller spores of *A. alternata.* A shorter lifespan of larger spores is unintuitive, as larger spores are often assumed to be more resilient than smaller spores (12; 43; 72; 73). Other species of *Alternaria* with large conidia also experience rapid declines in germinability when exposed to atmospheric conditions: in one experiment, 95% of *A. macrocarpa* conidia were unable to germinate after four days (10; 39). *Alternaria* fungi with large conidia are clustered in the monophyletic section *Porri* (see Figure 19 in 45). Perhaps species with large conidia are not under selective pressures to endure long-haul atmospheric travel because they settle out of the atmosphere quickly. At least among fungal spores that disperse in the atmosphere, we hypothesize a negative correlation between spore size and survival time in the atmosphere.

The timing of spore liberation will also influence effective dispersal. Lagomarsino Oneto et al. (5) establish the timing of spore ejection as playing a major role in determining how long spores dwell in the atmosphere before returning to the ground. For example, solar heat transfer causes atmospheric mixing, and consequently, all else being equal, spores released during the day settle less readily and will undergo longer journeys than spores released at night (5). Thus, we hypothesize that spore size, longevity, and the timing of spore release evolve and influence each other dynamically: spores undergoing long journeys facilitated by their small size and/or release patterns are selected for increased atmospheric survival, whereas spores traveling short distances resulting from their large size and/or release times in calm atmospheric conditions (e.g., at night), are under less selective pressure for longer-term atmospheric survival.

Our survival data are in broad agreement with data generated by other studies (11; 74; 75; 76; 77). For example, Magan et al. (68) also found RH as crucial to determining *A. alternata* germinability. Rotem et al. (10) found the germinability of *A. solani* conidia to decrease by 20% after eight hours’ exposure to sunlight. While UVB is clearly detrimental to germinability (Figure 3; Figure S2; Figure S3) in our experiments, interestingly, a small number of conidia exposed to UVB still germinated. We hypothesize these spores were shielded within clusters of spores. The clumping of dispersing spores is a rarely investigated phenomenon but it may be an important strategy used by fungi to survive harsh environments (4; 35; 78; 79; 80).

Tests of our hypotheses should include multiple large sets of closely related species with distinct spore morphologies and dispersal strategies. Because the genus *Alternaria* encompasses a diversity of spore shapes and sizes, it emerges as a model for studying the biophysical constraints and evolutionary tradeoffs of fungal dispersal within a phylogenetic framework.

Supporting Information can be found **here**

## Acknowledgments

We gratefully acknowledge support from UW-Madison’s Botany Department and funding from the United States Department of Agriculture, National Institute of Food and Agriculture, Hatch 1013478. In addition**, J.G.** was funded by a National Science Foundation Graduate Research Fellowship, North American Mycological Association Memorial Fellowship, and a Botany Department E.K. and O.N. **A.S.** and **D.L.O.** were funded by the European Research Council (ERC) under the European Union’s Horizon 2020 research and innovation programme (grant agreement No. 101002724 RIDING); the Air Force Office of Scientific Research under award number FA8655-20-1-7028. Allen Fellowship. **T.A.R.** was funded by the Genomic Sciences Program, U.S. Department of Energy, Office of Science, Biological and Environmental Research, as part of the Plant-Microbe Interfaces Scientific Focus Area at ORNL (http://pmi.ornl.gov); Oak Ridge National Laboratory is managed by UT-Battelle, LLC, for the U.S. Department of Energy under contract DEAC05-00OR22725. We are also grateful to Cécile Ané and Andrea Mazzino for their expertise and guidance throughout, and to Doug Sykes for making this study possible.

## Data Availability Statement

All data and scripts can be found at https://github.com/jacobgolan/Alternaria_Dispersal.git

## References

1. Bowden J, Gregory PH, Johnson CG. Possible wind transport of coffee leaf rust across the Atlantic Ocean. Nature. 1971 229(5285):500–1.

2. Purdy LH, Krupa SV, Dean JL. Introduction of sugarcane rust into the Americas and its spread to Florida. Plant disease (USA). 1985

3. Brown JKM, Hovmøller MS. Aerial dispersal of pathogens on the global and continental scales and its impact on plant disease. Science. 2002 297(5581):537–41.

4. Golan JJ, Pringle A. Long-Distance Dispersal of Fungi. Microbiol Spectr. 2017 5(4).

5. Lagomarsino Oneto D, Golan J, Mazzino A, Pringle A, Seminara A. Timing of fungal spore release dictates survival during atmospheric transport. PNAS. 2020 117(10):5134–43.

6. Malloch D, Blackwell M. Dispersal of fungal diaspores. In: The Fungal Community. New York: Marcel Dekker; 1992. p. 147–71.

7. Peay KG, Bruns TD. Spore dispersal of basidiomycete fungi at the landscape scale is driven by stochastic and deterministic processes and generates variability in plant–fungal interactions. New Phytologist. 2014; 204(1):180–91.

8. Aylor D. Aerial Dispersal of Pollen and Spores. The American Phytopathological Society; 2017. p. 418. (Epidemiology).

9. Maddison AC, Manners JG. Sunlight and viability of cereal rust uredospores. Transactions of the British Mycological Society. 1972 59(3):429–43.

10. Rotem J, Wooding B, Aylor DE. The role of solar radiation, especially ultraviolet, in the mortality of fungal spores. Phytopathology (USA). 1985.

11. Aylor DE, Sanogo S. Germinability of *Venturia inaequalis* conidia exposed to sunlight. Phytopathology. 1997 87(6):628–33.

12. Norros V, Karhu E, Nordén J, Vähätalo AV, Ovaskainen O. Spore sensitivity to sunlight and freezing can restrict dispersal in wood-decay fungi. 2015 5(16):3312–26.

13. Hawker LE, Madelin MF. The dormant spore. In: Weber DJ, Hess WM, editors. The Fungal Spore: Form and Function. New York: John Wiley; 1976. p. 235–44.

14. Hoekstra FA. Pollen and Spores: Desiccation tolerance in pollen and the spores of lower plants and fungi. In: Black M, Pritchard HW, editors. Desiccation and Survival in Plants: Drying Without Dying. CABI; 2002.

15. Ayerst G. The effects of moisture and temperature on growth and spore germination in some fungi. Journal of Stored Products Research. 1969 5(2):127–41.

16. Gervais P, Fasquel J-P, Molin P. Water relations of fungal spore germination. Appl Microbiol Biotechnol. 1988 29(6):586–92.

17. Magan N. Effects of water potential and temperature on spore germination and germ-tube growth in vitro and on straw leaf sheaths. Transactions of the British Mycological Society. 1988 90(1):97–107.

18. Piepenbring M, Hagedorn G, Oberwinkler F. Spore Liberation and Dispersal in Smut Fungi. Botanica Acta. 1998 111(6):444–60.

19. Pasanen A-L, Pasanen P, Jantunen MJ, Kalliokoski P. Significance of air humidity and air velocity for fungal spore release into the air. Atmospheric Environment Part A General Topics. 1991 25(2):459–62.

20. Verant ML, Boyles JG, Waldrep W, Wibbelt G, Blehert DS. Temperature-dependent growth of *Geomyces destructans,* the fungus that causes bat white-nose syndrome. PLoS One. 2012 7(9).

21. Park S, Chen Z-Y, Chanda AK, Schneider RW, Hollier CA. Viability of *Phakopsora pachyrhizi* urediniospores under simulated southern Louisiana winter temperature conditions. Plant Disease. 2008 92(10):1456–62.

22. Bonde MR, Berner DK, Nester SE, Frederick RD. Effects of temperature on urediniospore germination, germ tube growth, and initiation of infection in soybean by *Phakopsora* isolates. Phytopathology. 2007 97(8):997–1003.

23. Isard SA, Dufault NS, Miles MR, Hartman GL, Russo JM, De Wolf ED, et al. The Effect of Solar Irradiance on the Mortality of *Phakopsora pachyrhizi* Urediniospores. Plant Dis. 2006 90(7):941–5.

24. Kwon-Chung KJ, Sugui JA. *Aspergillus fumigatus—*What makes the species a ubiquitous human fungal pathogen? PLoS Pathog. 2013 9(12).

25. Jung JH, Lee JE, Lee CH, Kim SS, Lee BU. Treatment of fungal bioaerosols by a high-temperature, short-time process in a continuous-flow system. Appl Environ Microbiol. 2009 75(9):2742–9.

26. Koller LR. Ultraviolet Radiation. 2nd Edition. John Wiley & Sons; 1965. p. 314.

27. Robinson N. Global solar and sky radiation and their main spectral regions. In: Tromp SW, editor. Medical Biometeorology. Amsterdam, the Netherlands: Elsevier; 1963. p. 55–71.

28. Diffey BL. Solar ultraviolet radiation effects on biological systems. Phys Med Biol. 1991 36(3):299–328.

29. Iqbal M. An Introduction to Solar Radiation. Saint Louis, USA: Elsevier Science & Technology; 1983.

30. Zerefos CS, Bais AF. Solar ultraviolet radiation: modelling, measurements and effects. Berlin/Heidelberg, Germany: Springer Berlin / Heidelberg; 1997.

31. International Agency for Research on Cancer. Solar and Ultraviolet Radiation. In: IARC Monographs on the Evaluation of Carcinogenic Risks to Humans. Geneva, Switzerland: World Health Organisation; 2012. p. 363.

32. Sarantopoulou E, Stefi A, Kollia Z, Palles D, Petrou PS, Bourkoula A, et al. Viability of Cladosporium herbarum spores under 157 nm laser and vacuum ultraviolet irradiation, low temperature (10 K) and vacuum. Journal of Applied Physics. 2014 116(10):104701.

33. Parnell M, Burt PJA, Wilson K. The influence of exposure to ultraviolet radiation in simulated sunlight on ascospores causing Black Sigatoka disease of banana and plantain. Int J Biometeorol. 1998 42(1):22–7.

34. Singaravelan N, Grishkan I, Beharav A, Wakamatsu K, Ito S, Nevo E. Adaptive melanin response of the soil fungus *Aspergillus niger* to UV radiation stress at “Evolution Canyon”, Mount Carmel, Israel. PLoS One. 2008 3(8).

35. Li X, Mo JY, Yang XB. Frequency distribution of soybean rust urediospore clumps collected from naturally infected kudzu leaves in Nanning, China. Poster presented at: National Soybean Rust Symposium; 2006; St. Louis, Missouri.

36. Li X, Yang X, Mo J, Guo T. Estimation of soybean rust uredospore terminal velocity, dry deposition, and the wet deposition associated with rainfall. Eur J Plant Pathol. 2008 123(4):3

37. Norros V, Rannik U, Hussein T, Petäjä T, Vesala T, Ovaskainen O. Do small spores disperse further than large spores? Ecology. 2014 95(6):1612–21.

38. Hussein T, Norros V, Hakala J, Petäjä T, Aalto PP, Rannik Ü, et al. Species traits and inertial deposition of fungal spores. Journal of Aerosol Science. 2013 61:81–98.

37. Rotem J, Aust HJ. The effect of ultraviolet and solar radiation and temperature on survival of fungal propagules. Journal of Phytopathology. 1991 133(1):76–84.

38. Isard SA, Barnes CW, Hambleton S, Ariatti A, Russo JM, Tenuta A, et al. Predicting Soybean Rust incursions into the North American continental interior using crop monitoring, spore trapping, and aerobiological modeling. Plant Dis. 2011 95(11):1346–57.

39. Jongejans E, Skarpaas O, Ferrari MJ, Long ES, Dauer JT, Schwarz CM, et al. A unifying gravity framework for dispersal. Theor Ecol. 2015 8(2):207–23.

40. Woo C, An C, Xu S, Yi S-M, Yamamoto N. Taxonomic diversity of fungi deposited from the atmosphere. The ISME Journal. 2018 12(8):2051–60.

41. Calhim S, Halme P, Petersen JH, Læssøe T, Bässler C, Heilmann-Clausen J. Fungal spore diversity reflects substrate-specific deposition challenges. Scientific Reports. 2018 8(1):1–9.

42. Rotem J. The Genus *Alternaria*: Biology, Epidemiology and Pathogenicity. APS Press, American Phytopathological Society; 1994. p. 344.

43. Woudenberg JHC, Groenewald JZ, Binder M, Crous PW. *Alternaria* redefined. Stud Mycol. 2013 75(1):171–212.

44. Barberán A, Ladau J, Leff JW, Pollard KS, Menninger HL, Dunn RR, et al. Continental-scale distributions of dust-associated bacteria and fungi. PNAS. 2015 112(18):5756–61.

45. Ding S, Meinholz K, Cleveland K, Jordan SA, Gevens AJ. Diversity and virulence of *Alternaria* spp. causing potato early blight and brown spot in Wisconsin. Phytopathology. 2019 109(3):436–45.

46. National Agriculture Statistics Service (NASS). Press Release: 09/12/2019: Potato Summary. United States Department of Agriculture. 2019 (September).

47. Singh RP, Hodson DP, Huerta-Espino J, Jin Y, Bhavani S, Njau P, et al. The Emergence of Ug99 Races of the Stem Rust Fungus is a Threat to World Wheat Production. Annu Rev Phytopathol. 2011 49(1):465–81.

48. Prussin AJ, Li Q, Malla R, Ross SD, Schmale DG. Monitoring the long-distance transport of *Fusarium graminearum* from field-scale sources of inoculum. Plant Disease. 2013 98(4):504–11.

49. Bashan Y, Levanony H, Or R. Wind dispersal of *Alternaria alternata*, a cause of leaf blight of cotton. Journal of Phytopathology. 1991 133(3):225–38.

50. McCartney HA, Schmechel D, Lacey ME. Aerodynamic diameter of conidia of *Alternaria* species. Plant Pathology. 1993 42(2):280–6.

51. Psheidt JW. Epidemiology and control of potato early blight, caused by *Alternaria solani*. University of Wisconsin-Madison; 1985.

52. Ding S, Rouse DI, Meinholz K, Gevens AJ. Aerial concentrations of pathogens causing early blight and brown spot within susceptible potato fields. Phytopathology. 2019 109(8):1425–32.

55. Crutcher, H.L. 1969. Temperature & humidity in the troposphere. In Rex 1969, 45–84 (3)

53. Blumthaler M, Ambach W, Rehwald W. Solar UV-A and UV-B radiation fluxes at two Alpine stations at different altitudes. Theor Appl Climatol. 1992 46(1):39–44.

54. Blumthaler M, Ambach W, Ellinger R. Increase in solar UV radiation with altitude. Journal of Photochemistry and Photobiology B: Biology. 1997 39(2):130–4.

58. Dvorkin AY, Steinberger EH. Modeling the altitude effect on solar UV radiation. Solar Energy. 1999 65(3):181–7.

59. Hardin JW, Hilbe JM. Generalized Linear Models and Extensions. Stata Press; 2018. 598

60. Bolker BM. Post-model-fitting procedures with *glmmTMB* models: diagnostics, inference, and model output. 2020.

61. Stein AF, Draxler RR, Rolph GD, Stunder BJB, Cohen MD, Ngan F. NOAA’s HYSPLIT atmospheric transport and dispersion modeling system. Bull Amer Meteor Soc. 2015 96(12):2059–77.

62. Mesinger F, DiMego G, Kalnay E, Mitchell K, Shafran PC, Ebisuzaki W, et al. North American regional reanalysis. Bulletin of the American Meteorological Society. 2006 87(3):343–60.

63. D. Savage, M. J. Barbetti, M. J. MacLeod, M. U. Salam, M. Renton. Timing of propagule release significantly alters the deposition area of resulting aerial dispersal. Divers. Distrib. 2010 16:288–299.

64. Beljaars ACM, Holtslag A. M. Flux parameterization over land surfaces for atmospheric models. J Appl Meteor. 1991 30(3):327–41.

65. Hijmans RJ. Introduction to the *geosphere* package (Version 1.2-19). 2011.

66. Buliung RN, Remmel TK. Open source, spatial analysis, and activity-travel behaviour research: capabilities of the *aspace* package. Journal of Geographical Systems. 2008 10(2):191–216.

67. Pebesma EJ, Bivand RJ. Classes and methods for spatial data in R. R News. 2005 5(2):9–13.

68. Magan N, Cayley GR, Lacey J. Effect of water activity and temperature on mycotoxin production by *Alternaria alternata* in culture and on wheat grain. Appl Environ Microbiol. 1984 47(5):1113–7.

69. Pringle A. Asthma and the diversity of fungal spores in air. 2013 9(6):e1003371.

70. Bush RK, Prochnau JJ. Alternaria-induced asthma. Journal of Allergy and Clinical Immunology. 2004 113(2):227–34.

71. Pringle A, Baker DM, Platt JL, Wares JP, Latgé JP, Taylor JW. Cryptic speciation in the cosmopolitan and clonal human pathogenic fungus *Aspergillus fumigatus*. Evolution. 2005 59(9):1886–99.

72. Kauserud H, Colman JE, Ryvarden L. Relationship between basidiospore size, shape and life history characteristics: a comparison of polypores. Fungal Ecology. 2008 1(1):19–23.

73. Jones A.M. and Harrison R.M., The effects of meteorological factors on atmospheric bioaerosol concentrations–a review Review Sci Total Environ. 2014 326:151

74. Leach CM. Interaction of near-ultraviolet light and temperature on sporulation of the fungi *Alternaria, Cercosporella, Fusarium, Helminthosporium,* and *Stemphylium*. Can J Bot. 1967 45(11):1999–2016.

75. Fourtouni A, Manetas Y, Christias C. Effects of UV-B radiation on growth, pigmentation, and spore production in the phytopathogenic fungus *Alternaria solani*. Can J Bot. 1998 76(12):2093–9.

76. Braga GUL, Rangel DEN, Fernandes ÉKK, Flint SD, Roberts DW. Molecular and physiological effects of environmental UV radiation on fungal conidia. Curr Genet. 2015 61(3):405–25.

77. García-Cela ME, Marín S, Reyes M, Sanchis V, Ramos AJ. Conidia survival of *Aspergillus* section *Nigri, Flavi* and *Circumdati* under UV-A and UV-B radiation with cycling temperature/light regime. J Sci Food Agric. 2016 96(6):2249–56.

78. Dias, A. Epidemiological studies of shading effects on Asian soybean rust. Iowa State; 2008.

79. Furukawa S, Narisawa N, Watanabe T, Kawarai T, Myozen K, Okazaki S, et al. Formation of the spore clumps during heat treatment increases the heat resistance of bacterial spores. Int J Food Microbiol. 2005 102(1):107–11.

80. Schwinghamer EA. The relation of survival to radiation dose in rust fungi. Radiat Res. 1958 8(4):329–43.

